# Probing the structure-function relationship with neural networks constructed by solving a system of linear equations

**DOI:** 10.1101/2020.04.20.051565

**Authors:** Camilo J. Mininni, B. Silvano Zanutto

**Author notes:** Corresponding author (CJM).

## Abstract

Neural network models are an invaluable tool to understand brain function, since they allow to connect the cellular and circuit levels with behaviour. Neural networks usually comprise a huge number of parameters, which must be chosen carefully such that networks reproduce anatomical, behavioural and neurophysiological data. These parameters are usually fitted with off-the-shelf optimization algorithms that iteratively change network parameters and simulate the network to evaluate the changes and improve fitting. Here we propose to invert the fitting process by proceeding from the network dynamics towards network parameters. Firing state transitions are chosen according to the transition graph followed by an agent when solving a given behavioural task. Then, a system of linear equations is constructed from the network firing states and membrane potentials, in such a way that system consistency in guarantee. This allows to uncouple the activity features of the model, like its neurons firing rate and correlation, from the connectivity features and from the task-solving algorithm implemented by the network, allowing to fit these three levels separately. We employed the method to probe the structure-function relationship in a stimuli sequence memory task, finding solution networks where commonly employed optimization algorithms failed. The constructed networks showed reciprocity and correlated firing patterns that recapitulated experimental observations. We argue that the proposed method is a complementary and needed alternative to the way neural networks are constructed to model brain function.

## Introduction

Understanding brain function requires construction of physiological models that explain experimental data, which encompass behavioural outcome, anatomical features, neurons biophysics and coding properties, among others^1,2^. Many kinds of physiological models have been proposed along history, each one with their own merits. Among them, neural network models are well poised to connect all levels of analysis, from the behavioural to the molecular level, being a natural choice as neurons are the functional units of the brain. Yet, constructing neural networks that are suitable models is not an easy task. Neural networks can be hand-designed, setting network parameters following experimental data, or randomly chosen when experimental data is not available or as a mean of attaining more general conclusions. However, this approach may fall short given the complexity of the nervous systems. To tackle this issue, theorist have employed optimization methods to define the network parameters in such a way that a loss function is minimized. The loss function must encompass relevant aspects of the model, like its performance in one or several tasks, structural constrains such as the Dale’s principle, or a connectivity with a certain degree of sparseness ^3^. Optimization methods are widely used in artificial intelligence (AI), and the ongoing deep learning revolution has prompted an explosion of fitting algorithms, and the eagerness of taking advantage of them to build models of brain function^4,5^. However, AI needs are different from the theoretical neuroscience needs. Artificial intelligence deals with constructing systems capable of solving difficult tasks, employing very general optimization algorithms parameter fitting^6^. On the other hand, models in neuroscience are expected to explain how animals behave in simple tasks, yet with biologically plausible neural networks. Simple tasks are desired because behavioural outcome is easier to interpret, and mechanistic explanations easer to envisage. Thus, in AI the difficulty strives in the task, while in theoretical neuroscience it strives in the restrictions in network design that are imposed by biology. Therefore, methods for parameter fitting in theoretical neuroscience can take leverage from this point – the simplicity of the task – to solve problems that could be too hard to solve with generic optimization methods.

One approximation that has been overlooked consists in finding the synaptic weights of a network as the solution of a system of equations. For many commonly employed neural network models, neurons perform a weighted sum of their inputs, followed by a non-linear transformation. For these models, if neurons firing and their added postsynaptic potential are known, the synaptic weights can be readily found by solving a linear system of equations in which the neurons firing constitute the coefficient matrix and the added postsynaptic potentials are the dependent variables. Thus, the problem of finding the network parameters is replaced by the problem of finding sequences of valid network states that are consistent with solving the task. Although this problem might seem as hard as the former, we show in this work that viable network dynamics can easily be found by taking into account the transition graph associated with solving the task. By doing so, we were able to construct networks with millions of parameters extremely fast, without inefficient searches in parameter space. Moreover, optimization algorithms may have biases for a subset of all possible solutions^7^. These biases depend on the algorithms employed, the hyperparameters and the regularizations, and the relation between biases and its causes might be difficult to understand or control^8^. In contrast, our method allows to construct networks by sampling from a desired distribution of network dynamics, while further structural constrains on solutions can be easily imposed. Since the method proceeds from the network firing states to the network parameters, we call it the *Firing to Parameter* (FTP) method.

In this work we test the FTP method in a sequence memory task, and compare the method performance against an off-the-shelf optimization algorithm. Then, we show how to construct networks with certain activity and structural constrains, and analyse the relationship between structure and function.

## Results

### Neural networks that follow a predefined transition graph

We will consider an agent that interacts with its environment. At each time step *t* the agent is at one of *M* possible states *m*. Conversely, the environment adopts one of *L* possible states *e*. Agent and environment transitions can be expressed as:

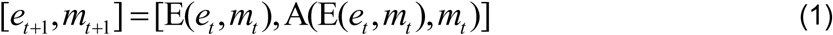

where E and A are the transition functions that take the agent and environment states and give the agent and environment states in the subsequent time step. The agent state *m* may codify several sub-states related to a biological agent, such as the behavioural response, reward signals, etc. Equation (1) thus describes a state machine which can model animal behaviour and neurophysiology. In particular, a behavioural task in which the agent must interact with the environment in a certain way to obtain reward can be codified in the E and A functions. Hence, any agent that solves a given behavioural task must follow the transition graph associated with solving that task. This includes agents controlled by recurrent neural networks, which are the main focus in this paper. We will work with networks of binary (McCulloch-Pitts) neurons composed of *N*_*z*_ recurrently connected integration neurons. Information about the environment is carried by a set of *N*_*y*_ sensory neurons (Fig. 1a). The temporal evolution of the network is dictated by:

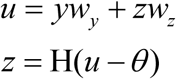

where *w*_*y*_ and *w*_*r*_ are synaptic weights matrices of sensory and integration neurons, vector *y* contains firing states of sensory neurons, *θ* is a vector of neuron thresholds, and H stands for the Heaviside function. Vector *u* contains the neurons activation states, akin of membrane potentials, and vector *z* contains the neurons firing states. We will call *z*_*m*_ the vector of firing states of all integration neurons associated with population state *m*, and *y*_*e*_ the vector of firing states of sensory neurons associated with environment state *e*. Since we want our network to follow the state transitions depicted in eq. (1), the next equation must hold:

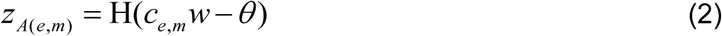

where *c*_*e,m*_ = [*y*_*e*_, *z*_*m*_] is the concatenation of sensory and integration neurons firing states, and *w* = [*w*_*y*_, *w*_*z*_] is the concatenation of the synaptic weight matrices. Equation (2) says that transitions in network states as ruled by the weight matrix *w* must be consistent with transitions in the transition graph that solves the target task.

**Figure 1.**
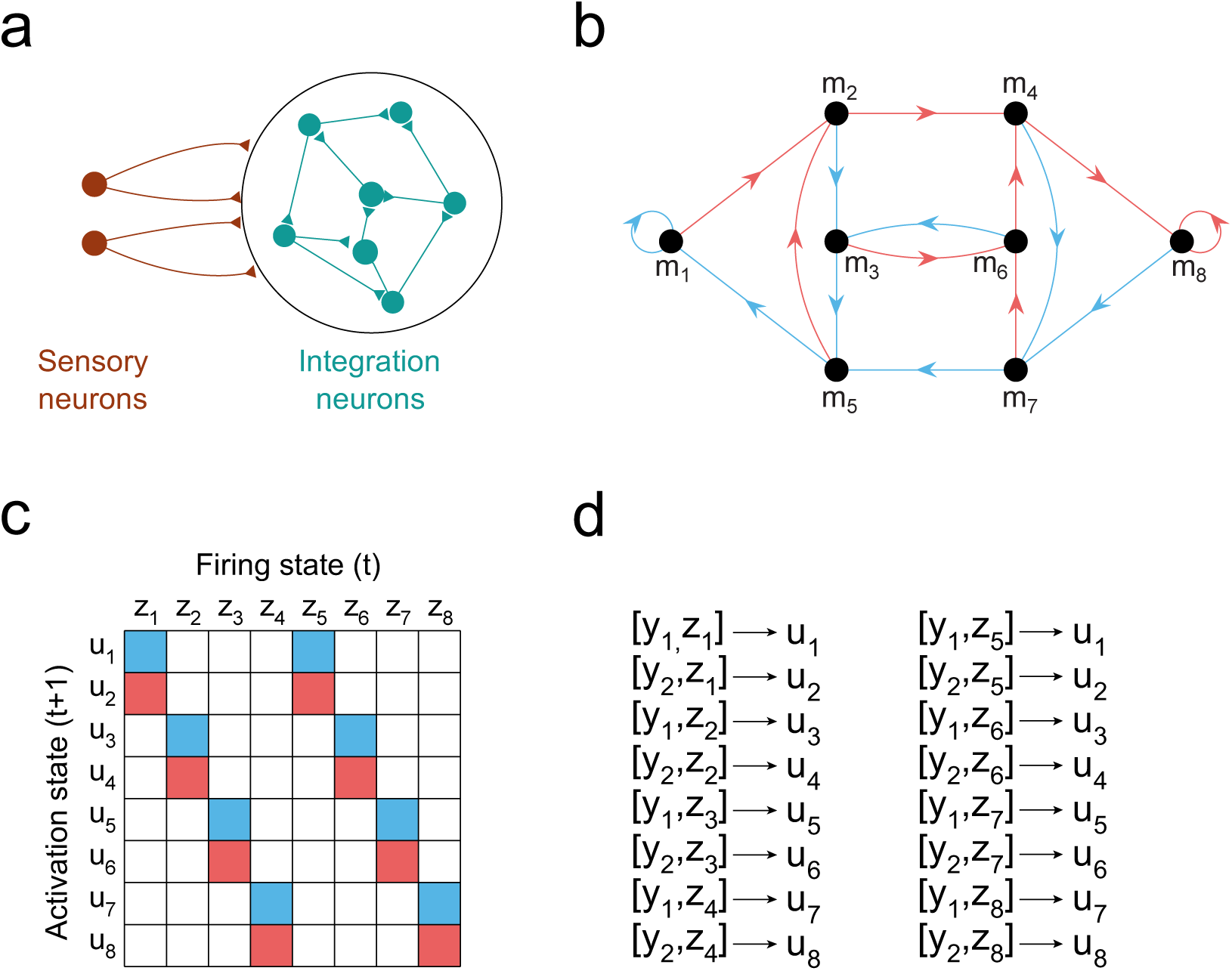
Constructing recurrent networks that follow a predefined transition graph. **a**, networks are composed of binary McCulloch & Pitts neurons. Sensory neurons codify external stimuli and project to the integration neurons through synaptic weights *w*_*y*_. Integration neurons are recurrently connected through synaptic weights *w*_*r*_. In the s-task, the population of integration neurons must codify in its firing state the sequence of the last *τ* stimuli presented. **b**, transition graph showing transitions between network states during execution of the s-task, for *τ* = 3. Each node in the graph is a network state, and arrows depict transitions between nodes after stimuli presentation. Each possible sequence of 3 stimuli in length is codified by exactly one network state. Nodes are numbered such that transitions can be represented in a simple transition matrix. **c**, transition matrix associated with the transition graph in panel (b). It shows the activation states *u* that are reached when recurrent neurons are in a population firing state *z*, and *s*_1_ (blue) or *s*_2_ (red) are presented. **d**, same transitions depicted in panels (b) and (c), but explicitly showing vectors *u*_*i*_ and vectors *c*_*i*_, which are the concatenation of one *y* and one *z*. The index *i* is such that *z*_*i*_ and *u*_*i*_ are the firing state and activity vectors corresponding to agent state *m*_*i*_.

### Linear system construction

We are interested in constructing a recurrent network of neurons by solving a system of linear equations built from the set of neurons firing states. To this end we first define a coefficient matrix *C*, whose rows result from the concatenation of one *y*_*e*_ vector and one *z*_*m*_ vector, for all combinations of environment state *e* and agent state *m*:

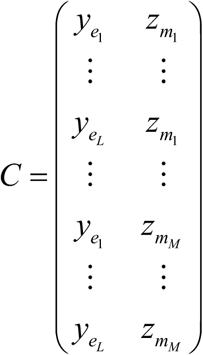

Then we define a matrix *U*, such that its ith row vector *u*_*i*_ is such that H(*u*_*i*_ −*θ*) gives the *z* associated with the state *m* that should follow from the *e* state and *m* state of row *i* in *C*, according to the desired transition graph. Thus, matrix *C* and *U* condense all the transitions required to solve the target task. It follows that:

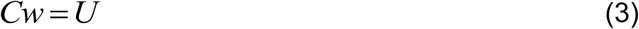

The matrix *w* can be found by computing:

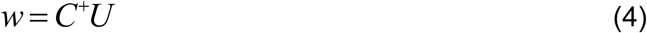

where *C*^+^ stands for the pseudoinverse of *C*. Thus, the connectivity matrix *w* can be obtained by solving the linear system with coefficient matrix *C* and dependent variable matrix *U*. Since we are using the pseudoinverse to solve the system, the solution is the one with minimum Frobenius norm^9^.

We want to sample from the set of matrixes *w* that accomplish the task constrain, i.e. networks constructed with the sole constrain of solving the target task. A naïve approximation to this problem would be to pick the *u*_*i*_ vectors at random, threshold them to obtain the associated *z*_*i*_ vectors, and construct matrices *C* and *U* by following the desired transition graph. However, by doing so it is very likely that we end up by having an inconsistent system of equations, meaning that there is no network of neurons that can follow those state transitions. This is because matrix *C* is not full rank, but its rows are linearly dependent. If we consider the case of two stimuli *s*_1_ and *s*_2_, codified by vectors *y*_1_ and *y*_2_, then each vector [*y*_1_, *z*_*m*_] can be expressed as a linear combination of [*y*_2_, *z*_*m*_] and vectors [*y*_1_, *z*_*P*_] and [*y*_2_, *z*_*P*_], where vector *z*_*P*_ can be any vector taken from the set of all firing states the network can adopt:

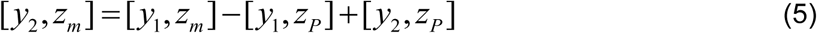

Thus, rank(*C*) = *M* +1. Following the Rouché-Capelli theorem^10^, eq. (3) has a solution if and only if rank(*C*) = rank([*C,U*]), being [*C,U*] the augmented matrix. Yet, if we choose vectors *u* randomly, when adjoined to matrix *C* the linear dependencies expressed in eq. (5) will be broken, and the resulting augmented matrix will have rank above *M* +1. However, consistency can be enforced if initial randomly generated vectors *u* are linearly combined following linear dependencies in *C*, such that the same linear dependencies in *C* are conserved in the augmented matrix.

### The s-task

In the following we will consider a sequence memory task: the environment consists of two stimuli *s*_1_ and *s*_2_, which are sequentially presented at each time step, chosen randomly with equal probability. To obtain reward at time step *t* the agent has to recall the stimulus presented at time step *t* − Δ*t*. Successful behaviour thus requires to have a memory of stimuli sequences of length *τ* = Δ*t* +1, starting from *t* − Δ*t*. The constant *τ* defines the memory requirements of the task. Figure 1b shows an agent’s states and the transitions gated by the stimuli when solving the s-task for *τ* = 3.

To solve the task the agent needs at least *M* = 2^*τ*^ states, meaning that complexity grows exponentially. This would suggest that the task is a complex one. However, it can be seen that the transition matrix has a stereotyped form if nodes are numbered properly (Fig. 1c). The transition graph in Fig. 1b shows the state transitions any agent that solves the s-task should follow. With the transition matrix structure at hand we can construct matrices *C* and *U*. We will define a neural network with two sensory neurons such that 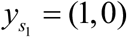 and 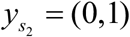. If we order the transitions as in Fig. 1b-d, we have that, if eq. (3) has a solution, rows in the augmented matrix should satisfy:

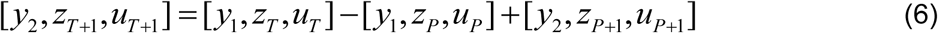

where *T* are indexes over the rows of *C* and *U*, which are odd numbers between 1 and *M*. The row index *P* is and odd number between 1 and *M*, different from all *T*. Note that *z*_*T*_ = *z*_*T* +1_ and *z*_*P*_ = *z*_*P*+1_, but *u*_*T*_ ≠ *u*_*T* +1_ and *u*_*P*_ ≠ *u*_*P*+1_. Equation (6) shows us how row vectors in matrix *U* should be linearly combined such that eq. (3) has a solution. We have that:

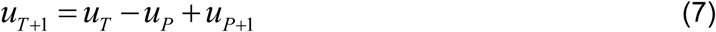

This means that the number of linear combinations in *U* is *R* = 2^*τ*^ / 2 −1, and rank(*U*) = 2^*τ*^ / 2 +1. Note that rewriting eq. (7) we have:

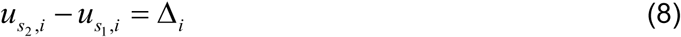

Where 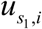 and 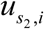 are the activation that neuron *i* adopts after presentation of *s*_1_ and *s*_2_, respectively. In words, eq. (8) tells that the difference in effects provoked by the stimuli is a constant for each neuron, regardless of which network state or transition we are dealing with. This fact is not surprising, since synaptic weights are held fixed, so each stimulus has the same effect at any time, which is specific for each neuron. Thus, making the system of equations in (3) consistent is equivalent to guarantee that activation values are chosen so that the effect of each stimulus is consistent.

We proceeded by generating a vector of thresholds *θ*, with 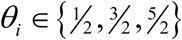. Then, we constructed base matrix *U*_*base*_ with *M* _*base*_ = 2^*τ*^ / 2 +1 row vectors such that:

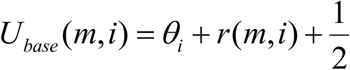

Where *r*(*m,i*) is an integer uniformly sampled from the [-5,5] interval. We added the term 1_2_ to avoid fitting errors when numerically solving the system, otherwise the activation values could be equal to the threshold values, which would result in erroneous firing states because of numeric precision issues. This initial randomly generated matrix *U*_*base*_ has full rank. We computed the vector Δ of Δ_*I*_ elements as the difference between the first two rows of *U*_*base*_. Next, we applied eq. (8) to generate the remaining *R* rows as linear combinations of the third to the last row of *U*_*base*_, obtaining 2^*τ*^ row vectors which constitute the matrix *U*^*^. Each row vector *u* in matrix *U*^*^ has the neurons activations for one of the 2^*τ*^ network state. Applying eq. (8) creates a dependency between 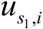 and 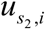. Hence, for each linear combination we chose at random which activation value (the one associated with *s*_1_ or *s*_2_) will be defined in terms of the other. This is to ensure that *u* value distributions are equal between stimuli. We constructed matrix *Z* by applying threshold *θ* to *U*^*^, and then we followed the ordering depicted in Fig. 1b-d to construct matrix *U* from *U*^*^, and matrix *C* from *Z* and vectors *y*_1_, *y*_2_. Finally, we employed eq. (4) to obtain the synaptic weight matrix *w*. Since matrix *w* is the minimum Frobenius norm solution to eq. (4) and defines a network that solves the s-task, we call it a *T+F* network.

We are assuming that two conditions are met after thresholding: 1) the resulting vectors *c* are all different, and 2) they are linearly independent. If after thresholding any vector is repeated, this would result in lower performance in the task, since not all sequences of length *τ* will be encoded. On the other hand, if linear independency fails after thresholding, then matrix *C* will have more linear combinations than the contemplated in eq. (5), meaning that combining the rows of *U* following eq. (8) will not be enough, and some linear dependencies in *C* will be lost in the augmented matrix, making the system inconsistent. In our implementation of the algorithm, if any of these two conditions were not verified, then the algorithm was restarted from the beginning. This occurred sometimes, for *τ* < 5. For higher *τ*, both conditions were always fulfilled in one attempt.

In the above explanation we assumed that *N* _*z*_ = 2^*τ*^, such that there is one neuron per sequence of length *τ*. It was possible to fit networks with lower number of neurons, but undesired linear dependencies in *C* after thresholding, or a number of network states bellow 2^*τ*^ occurred with higher probability, especially for *τ* > 3.

We employed the FTP algorithm to construct networks of *N* _*z*_ = 2^*τ*^ neurons that solve the s-task (Fig. 2a,b). The resulting synaptic weight distribution had zero mean and resembled a normal distribution, at least for the *w*_*r*_ values (Fig. 2c). In fact, the synaptic weight distributions became progressively closer to a normal distribution as more neurons were used in network construction (Fig. 2d). We also noted that the absolute weight value decreased, especially for *w*_*r*_ values (Fig. 2e), which can be explained by thinking that more neurons imply more parameters and hence more degrees of freedom to reach a lower Frobenius norm. This observation will become important later when imposing structural constrains to the network.

**Figure 2.**
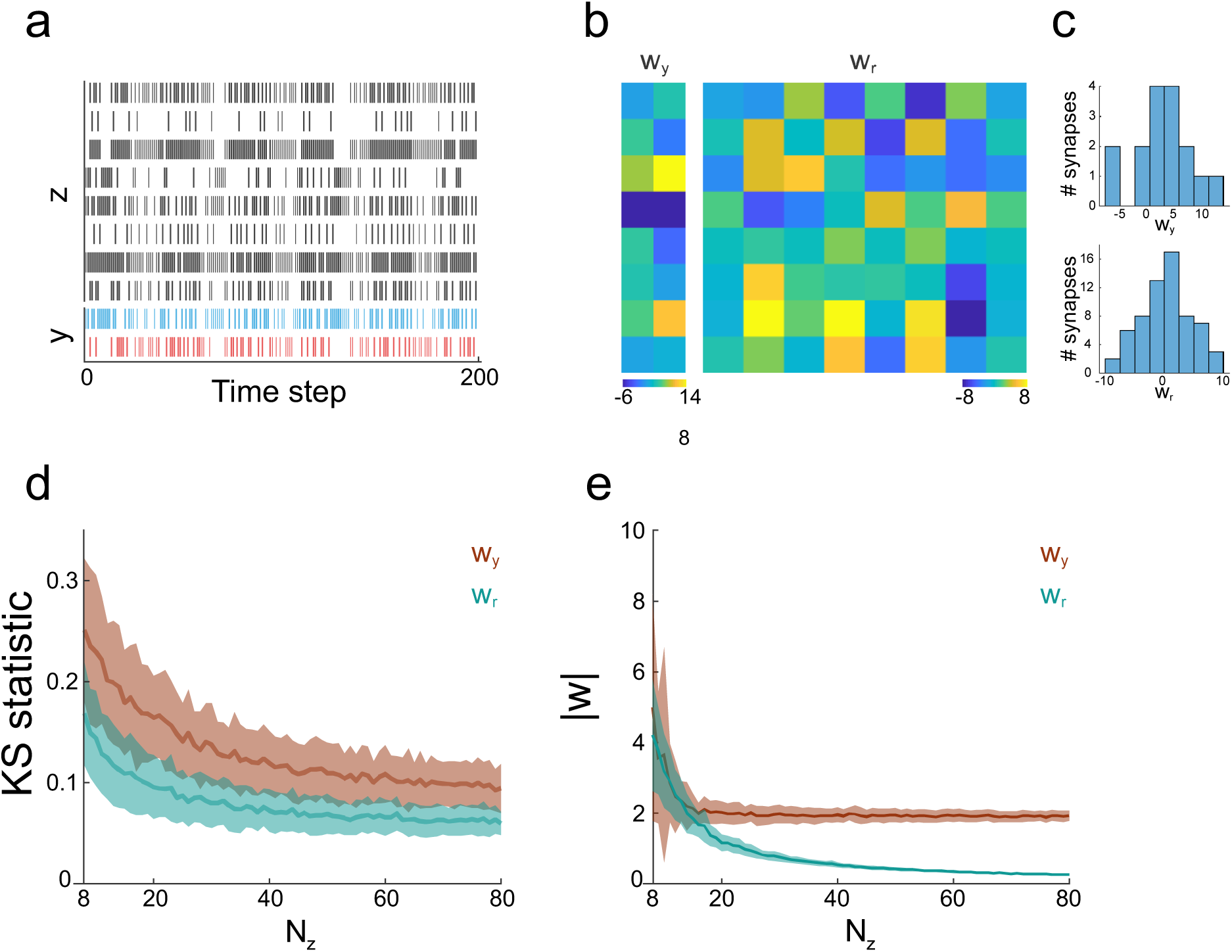
Solving the s-task with the FTP algorithm. **a**, raster plot showing the neurons firing states in a network constructed to solve the s-task for *τ* = 3. The network is composed of eight integration neurons and 2 sensory neurons. Each possible sequence of 3 stimuli has a unique network firing state that codifies it. Therefore, the network has 8 possible firing states. **b**, sensory and integration synaptic weights of the network. For this example (*f*_*r*_ = 1) the range of weights is grater for sensory than for integration synapses. **c**, distribution of synaptic weights for sensory (upper panel) and integration (lower panel) synaptic weights, for the same network as in (a, b). Distributions are zero centred. **d**, Kolmogorov-Smirnov statistic between the distribution of synaptic weights and a normal distribution of the same mean and variance. As the number of integration neurons increases, the distribution of synaptic weights gets closer to a normal. Integration neurons weights are closer to a normal than sensory neurons weights. Mean ± SD are shown for n = 100 networks that solves the s-task, with *τ* = 3. **e**, absolute synaptic weight values as a function of the number of integration neurons in the network. Absolute values are higher and of larger variability when the neuron count is close to the number of coded stimuli sequences. As the number of integration neurons increases the absolute mean value and dispersion decreases. Sensory neurons weights quickly reach a minimum, while integration neurons weights decrease in the entire range of integration neurons. Mean ± SD are shown for n = 100 networks that solves the s-task, with *τ* = 3.

### Efficiency of the FTP algorithm

We assessed the performance of the algorithm by measuring the time expended in finding solutions for *τ* = 1 to *τ* = 12, and comparing these times with the times required for a genetic algorithm (GA) to find solutions for the same *τ* values and number of neurons. The FTP outperformed the genetic algorithm by several orders of magnitude (Fig. 3a), and for *τ* > 6 the GA could not find a network with performance above 0.9. The result is not surprising, since the time complexity of solving a linear system of equations is *O*(*n*^3^) ^10^, with *n* being the total number of variables. As *τ* was increased, most of the GA running time was expended in the simulation of the networks, while the fraction of time expended in evaluating network performance with the classifier became smaller.

**Figure 3.**
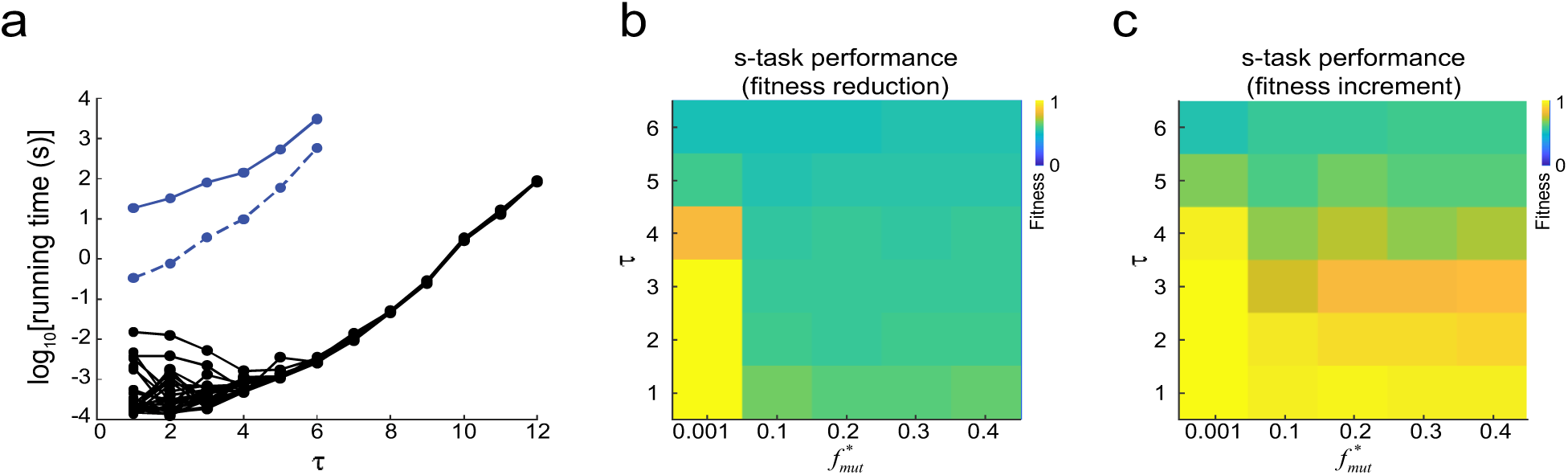
Efficiency of the FTP algorithm. **a**, efficiency of FTP and GA, measured as the time expended in finding a network solution for s-tasks of different *τ*. The time expended by the FTP algorithm is orders of magnitude lower than the time expended by the GA. As *τ* increases, the time expended in network simulation (dashed blue line) tends to match the total time expended by the GA (solid blue line). The FTP running time showed some variability for *τ* < 6, in the range of tens of milliseconds. The absence of points in the GA curves for *τ* > 6 means that this algorithm could not find a solution within the limit of 1 hour of running time. The GA was run 1 time for each *τ*, while the FTP was run 30 times for each *τ*. **b**, fitness of solution networks found for different *τ* after their synaptic weights were mutated for 20 generations with different mutation ratios, 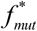 and selected to reduce their fitness. Mutations reduced fitness to chance level for all 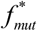 values except for the lowest. For *τ* ≥ 5 even the lowest 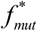 had detrimental effects. **c**, fitness measured after networks in (b) where subjected to 20 generations of mutation and selection to increase their fitness. Fitness could be restored when *τ* and 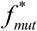 values were low. If *τ* or 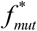 where higher, restoration was only partial or did not occurred.

Given that performance of optimization algorithms is sensitive to several hyperparameters, including initial conditions and mutation factor, we asked whether a GA that starts near a solution network would stay around the solution or would it drift away. To that end we employed the FTP to find a solution network, and set it as the individual from which the first population was built. Then the GA was used to reduce fitness over 20 generations, such that the population became 20 generations apart from the solution (Fig. 3b). Next, the GA was run for another 20 generations in the increasing fitness direction (Fig. 3c). In each generation, random mutations were applied, with a mutation factor 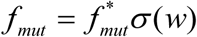 (see Methods). We run this experiment for several values of *τ* and 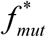. It can be seen that up to *τ* = 4 performance dropped to chance level during the fitness reduction phase, followed by a total or partial recovery during the fitness increment phase. However, performance did not recover for higher values of *τ*. This suggests that, as *τ* increases, the basin of attraction around the initial solution gets narrower, leading the GA to drift away from the solution. Although these results cannot rule out that other optimization algorithms have better performance, they do highlight how small the solution space is, and the huge gap between the FTP and another, more generic fitting algorithm.

### Imposing activity constrains through *U* matrix initialization

We want to construct neural network models that not only solve relevant tasks but do so under desired firing constrains, as measured in real brains. Some of these constrains are low firing rates (FR)^11,12^, or low correlation coefficient (CC)^13^. In regular optimization algorithms, these constrains can be imposed to solution networks by introducing regularization terms in the loss function^3^. On the other hand, in the FTP algorithm the activity states of the network are the result of linearly combining the rows of an initial matrix *U*_*base*_. Hence, we can apply firing constrains by appropriately choosing this initial matrix. For example, to attain networks that solve the s-task with low/high firing it suffices to choose an initial matrix *U*_*base*_ such that after thresholding the resulting matrix *C* has few/many ones. Following this procedure, we constructed networks with average FR within a wide range of target FR (Fig 4a, blue line). Shuffling the afferent synaptic weights of each neuron produces only small changes to the average FRs (Fig. 4a, red line). This suggests that it is the distribution of afferent synaptic weights the critical structural statistic that defines the networks average FR, and not its precise connectivity. Solutions are harder to find for extreme FR values, because thresholding gives *C* matrices with repeated rows, which translate in not enough network states to codify all stimuli sequences.

**Figure 4.**
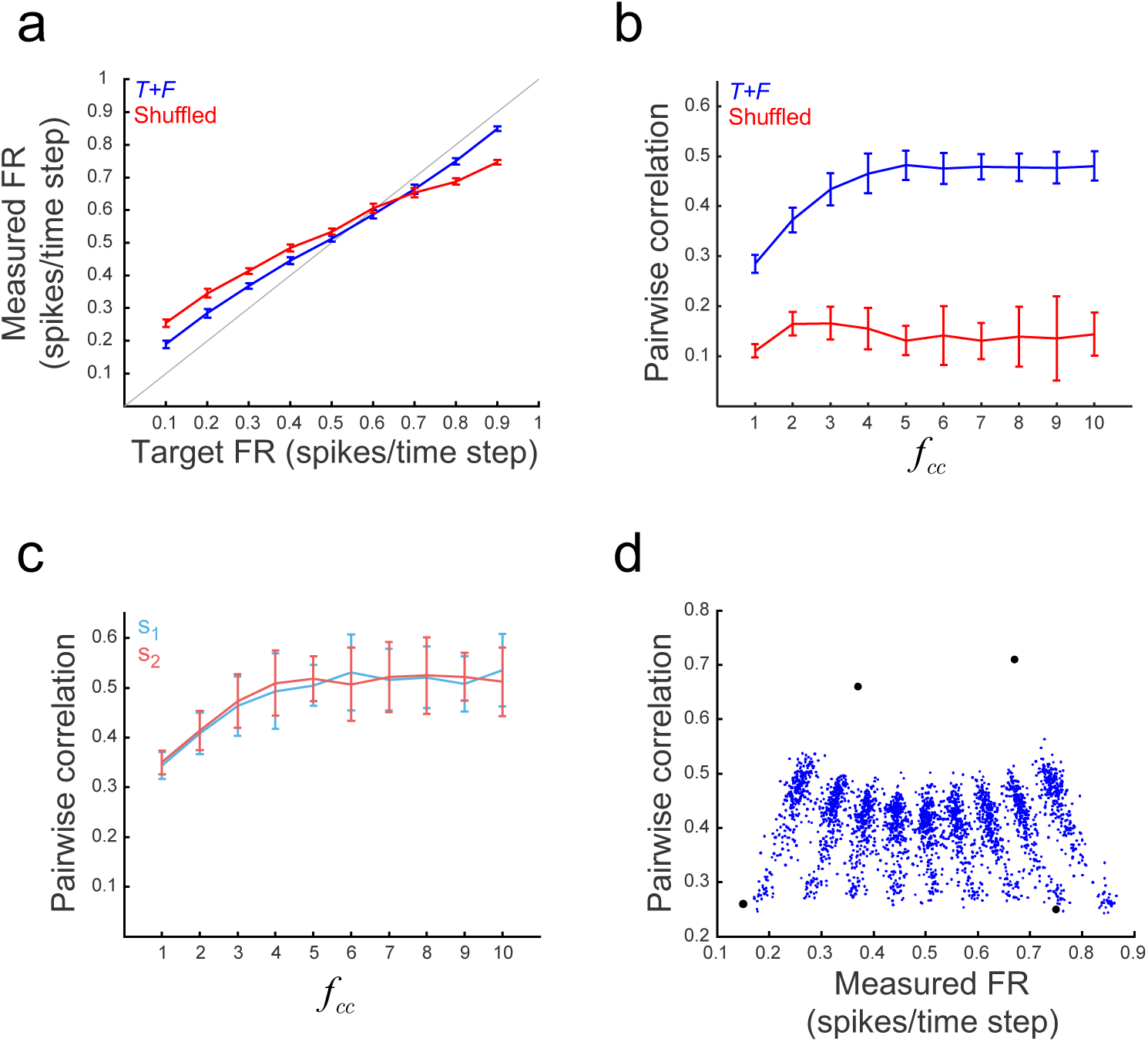
Using FTP to construct networks subjected to task and activity constrains. **a**, real FR measured in networks constructed to solve the s-task, as a function of the target FR. The FR of networks constructed with the FTP algorithm are close to the target FR (blue line). There is a tendency to obtain lower firing rates for target FR values above 0.5 spikes/ time step, and higher firing rates for target FR values bellow 0.5 spikes/time step (grey line is the identity function). The same networks with their afferent synaptic weights shuffled (red line) show a similar relationship between target FR and measured FR, albeit with a lower slope. A total of 30 networks were generated for each target FR. Mean ± SD are shown, n = 30. **b**, correlation between pairs of integration neurons as a function of the scaling factor *f*_*cc*_. Pairwise correlation, computed over all time steps, increases with *f*_*cc*_ until it saturates at CC = 0.48 for *f*_*cc*_ ≥ 5 (blue line). Networks with their afferent synaptic weights shuffled (red line) show low correlation, invariant to *f*_*cc*_. A total of 30 networks were constructed for each *f*_*cc*_ value, with target FR set to 0.1 spikes/time step. Mean ± SD are shown, n = 30. **c**, pairwise correlation computed separately for *s*_1_ and *s*_2_ (noise correlation). The correlation coefficient increases with *f*_*cc*_, similarly for both stimuli, and closely following correlation values in (b). Mean ± SD are shown, n = 30. **d**, Measured FR as a function of pairwise correlation. Each blue dot shows the FR and CC of one network constructed to solve the s-task with FR and correlation constrains imposed by *U*_*base*_ initialization. Values for 2700 networks are shown. Points form stripes pointing towards FR = 0.5 spikes/time step, each stripe corresponding to networks with the same target FR. As correlation increases, the measured FR tends to 0.5 spikes/time step. Black dots show FR and correlation of 4 networks for which FR and correlation constrains were imposed by evolution of a population of *U*_*base*_ matrices.

On the other hand, we can construct networks with desired signal correlation, by multiplying Δ by a factor *f*_*cc*_, which results in stimuli inducing different firing rates (Fig. 4b). For networks shown in Fig. 4 (*τ* = 4, *f*_*r*_ = 3) correlations could be modulated in a range between 0.25 and 0.5. Although scaling of Δ is expected to induce signal correlation, it can be seen that it is inducing noise correlation as well, as a by-product (Fig. 4c). Correlations of networks solving the s-task are significantly higher than correlations of their synaptic weights-shuffled counterparts (Fig 4b, blue vs. red), which shows that pairwise correlations depend on the whole weight matrix and not only on the distribution of the afferent weights, as is the case with FR. It also suggests that the set of networks that solve the s-task necessarily have correlation above a minimum. On the other hand, correlations also seem not to exceed a certain value: higher correlations would imply a reduced number of network states, incompatible with the number of sequences required to codify.

Hand-based manipulation of *U*_*base*_ allows to generate solution networks in a wide range of FR and CC (Fig. 4d, blue dots). An even better control of firing and correlation can be achieved by fitting *U*_*base*_ by means of a GA in which the fitness of *U*_*base*_ is a function of the FR and CC computed over the population firing states of the network generated from that *U*_*base*_. Fitting *U*_*base*_ allows for more extreme values of FR and CC (Fig 4d, black dots), while keeping computational cost low by computing FR and CC over the set of population firing vectors *c* instead of computing the actual network activity by simulating the network. Altogether, both methods (*U*_*base*_ manipulation, or its evolution with a GA) easily allow to generate networks that perfectly solve the task, while imposing desired activity constrains at the same time.

### Applying structural constrains with projected gradient descend in isofunction weight space

Networks generated so far share one structural constrain: their synaptic weights matrix is the one that minimizes the Frobenius norm. Other relevant structural constrains, such as the lack of self-connections, Dale’s principle, or sparse connectivity are not satisfied. Since these structural constrains are key experimentally observed features(Lefort et al. 2009; Seeman et al. 2018; Strata and Harvey 1999) but see^17,18^, we were interested in imposing such constrains onto the *w* obtained by the algorithm. To do this we followed a projected gradient descent (PGD) approach^19^, taking advantage of the fact that the loss function ℒ, which encloses the structural constrains, is a linear function with respect to the synaptic weights, and that the matrix *w* can be changed without changing the stimulus-response mapping (see Methods). To exemplify the procedure we constructed a network that solves the s-task for *τ* = 4, with *f*_*r*_ = 3 (Fig. 5a), and then we employed PGD to transform its matrix *w* to remove self-connections, enforce Dale’s principle with a 4:1 Ex:In ratio, and set a sparsity *sp* = 40% (defined as the percentage of weights equal to zero). The PGD reduced the loss function ℒ in a steady fashion, reaching a negligible error, provided that the network had enough neurons (Fig. 5b). It is remarkable how such different synaptic weight matrices, as the ones depicted in Fig. 5a,c,d, gave rise to exactly the same stimulus-response mapping.

**Figure 5.**
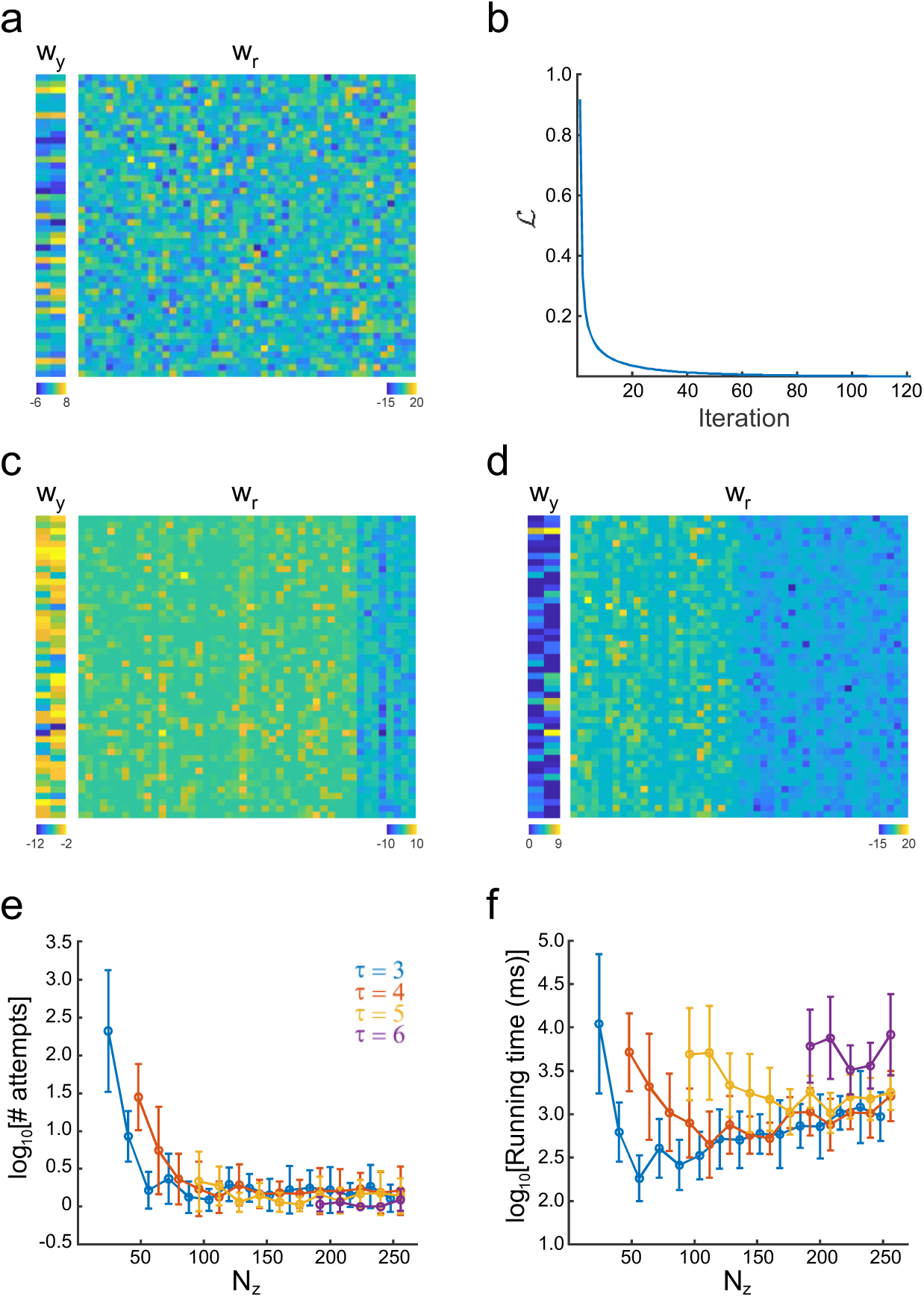
Applying structural constrains to networks. **a**, synaptic weight matrices *w*_*y*_ and *w*_*r*_ of the network obtained through FTP before structural constrains were imposed, The network was constructed to solve the s-task for *τ* = 4 and *f*_*r*_ = 3, with a target FR of 0.1 spikes/time step. **b**, Loss function ℒ as a function of the number of iterations of the PGD algorithm. The loss function falls below the criterium *e*_1_ = 10^−3^ at iteration 121. **c, d**, synaptic weight matrices *w*_*y*_ and *w*_*r*_ for a network with the same stimulus-response mapping but after applying structural constrains: (c) no self-connection, Dale’s principle, with 40 excitatory and 10 inhibitory neurons, and sparsity *sp* = 40%; (d) no self-connections, Dale’s principle, with 26 excitatory and 24 inhibitory neurons, and sparsity *sp* = 23%. **e**, average number of attempts to obtain one network with successful structural fitting, as a function of the number of integration neurons, and for different *τ*. The number of attempts is high when the neuron number is low, but it decreases fast as the neuron number increases. From 60 neurons onwards, less than five attempts are needed, on average, to obtain one network with the desired structural constrains. Mean ± SD are shown. **f**, total running time to obtain one network with successful structural fitting, as a function of the number of integration neurons, and for different *τ* (color code as in (g)). Running time decreases and then increases for *τ* = 3 and *τ* = 4. The case of *τ* = 6 is the one with more neurons and equations to solve, and present some of the highest running times, even when the number of neurons is high. Nevertheless, all average running times are below the tens of seconds.

We noted the that structural constrains could not be imposed to networks with low number of neurons, i.e. *N* = *N* _*y*_ + *N*_*z*_ between *M* and 3*M*. This is not surprising, since it is expected that imposing more constrains requires more parameters. To evaluate the efficiency of the PDG in relation with the number of neurons, we imposed the above structural constrains for networks solving the s-task with *τ* = 3 to *τ* = 6, and *N*_*z*_ between 32 and 256 neurons. Since matrix *U* and vector *θ* were randomly chosen, it is expected that some of them result in matrices *w* for which the structural constrains are impossible to apply. Consequently, we measured PDF efficiency by computing *# attempts*, the number of networks that were required to generate until obtaining the first successfully constrained network. It can be seen that *# attempts* decreased as the number of neurons increased (Fig. 5e). Concordantly, the computing time required to obtain a fitted network decreased as the number of neurons increased, because fewer fitting attempts were required (Fig. 5f). The fitting time was somewhat higher for networks with the highest neuron count, but always within the order of tens of seconds, even for *N*_*z*_ = 256.

### Linking structure, function and activity

Neural network structure determines its activity, which in turn translates into function. To understand the relationships between these three network features we constructed networks with different average firing rate and functionality and analysed their structure, more precisely, their connectivity. One key aspect of connectivity is reciprocity, which has been observed experimentally^20^ and its implications studied theoretically^21^. Here we chose the correlation between weights of incoming and outgoing synapses as the measure of reciprocity^22^ (see Methods).

We have already shown how the FTP algorithm can be employed to generate networks with predefined activity features, namely, with desired firing rate and correlation. To compare networks with different *functionality* we constructed networks which had the same number of neurons and network states but for which the graph of transitions between network states was generated at random (Fig. 6a,b). In this manner we can construct networks whose dynamics have complexity similar to that of networks that solve the s-task, but which lack their function, i.e. to codify sequences of stimuli of length *τ*. We screened networks with memory ranging from *τ* = 2 to *τ* = 7, and FR from 0.1 spikes/time step to 0.9 spikes/time steps, and found that the reciprocity varied with *τ*, FR and neuron number. In particular, we observed that, when *f*_*r*_ = 1, reciprocity was positive and of lower mean for networks that were the minimum Frobenius norm solution to the s-task (*T+F* networks, Fig. 6c), in comparison with networks that were the minimum Frobenius norm solution to a random transition graph (*F* networks, Fig. 6d). However, for bigger *f*_*r*_ the relationship was inverted, and *T+F* networks showed positive reciprocity (Fig. 6e) while *F* networks showed negative reciprocity (Fig. 6f).

**Figure 6.**
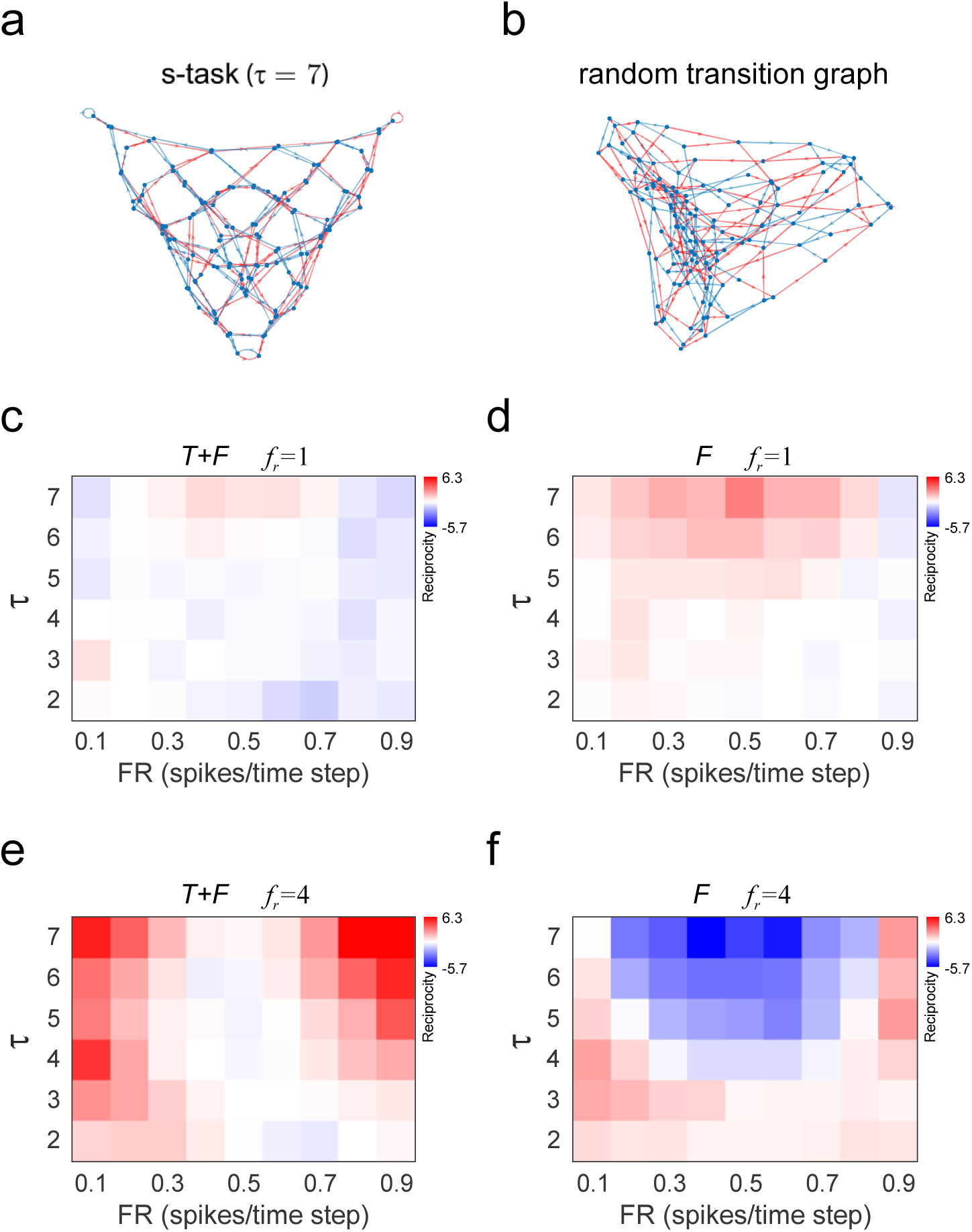
Reciprocity as a function of *τ*, FR and the type of transition graph. **a**, transition graph for solving the s-task with *τ* = 7. Blue and red lines represent transitions gated by *s*_1_ and *s*_2_, respectively. **b**, random transition graph. Nodes (network states) may receive different number of incoming connections. There are 24 nodes that are gated by both stimuli. **c**, reciprocity for *T+F* networks, with *f*_*r*_ = 1, as a function of *τ* and target FR. Reciprocity changes from slightly negative to slightly positive as *τ* increases. For *τ* = 7, reciprocity is maximized around target FR = 0.5 spikes/time step, and decreases for lower and higher values of target FR. **d**, *F* networks with *f*_*r*_ = 1 shows increasing positive reciprocity as *τ* increases, maximized at target FR = 0.5 spikes/time step. **e**, when the number of neurons is higher (*f*_*r*_ = 4), *T+F* networks show positive reciprocity that is minimal around target FR = 0.5, and increases towards higher and lower target FR, reaching the highest reciprocity values among all networks screened. **f**, reciprocity of *F* networks gets increasingly negative as *τ* increases, reaching the lowest reciprocity among all networks screened, around target FR = 0.5 spikes/time step. For all panels, 30 networks were constructed for each *τ* and target FR combination. Normalized means (mean/SD) are shown. Positive and negative reciprocity values were mapped separately to colours red and blue, respectively. Red tones go from 0 reciprocity (white) to maximal positive reciprocity (pure red). Blue tones go from 0 reciprocity (white) to maximal (in absolute value) negative reciprocity (pure blue). All random graphs were constructed with *f*_*bc*_ = 0.5. Graphs were plotted with the Force-directed layout

To further describe these relationships, we selected networks constructed for *τ* = 7 and *f*_*r*_ = 1 (Fig. 7a-c) and *f*_*r*_ = 4 (Fig. 7d-f). Signal and noise correlation varied with FR following an inverted U-shape relationship, with a maximum next to 0.5 spikes/time step (Fig. 7a,d). Note that, up to a FR = 0.5 spikes/time step, CC increased with FR, as has been observed experimentally^23,24^. Interestingly, the way CC changed with FR was similar for both *T+F* and *F* networks, with the distinction that *F* networks had overall higher correlations than *T+F* networks. We computed the CC after shuffling the inter-spike interval of each neuron and found that it remained practically invariant with respect to FR. These CC values were also much smaller than the CC values of the non-shuffled firings (CC_shuffled_ = 0.0224 ± 1.10^−4^, mean ± SD, for n = 90 *T+S* networks pooled over all FR values). These results rule out the possibility that correlations were trivially increasing with FR because of the higher number of spikes. We also found an inverted U-shape between reciprocity and FR, and a linear relationship between reciprocity and CC. The parabolic relationship is accentuated in networks with more neurons, the curve being more pronounce and of lower dispersion. With *f*_*r*_ = 1 reciprocity tended to be maximized as FR approached 0.5 spikes/time step, with *F* networks showing higher (and positive) reciprocity (Fig. 7b). With *f*_*r*_ = 4, reciprocity tended to increase as FR departed from 0.5 spikes/time step towards lower and higher values, i.e., networks with lower correlation (Fig. 7e). Specially, *F* networks showed negative reciprocity for all firing rates, except for the more extreme cases (0.1 and 0.9 spikes/time step). Just as the reciprocity/FR relationship inverts with the number of neurons, so does the reciprocity/CC relationship. Networks with higher reciprocity has higher correlation when the number of neurons is low (Fig. 7c). However, and somewhat counterintuitive, when the number of neurons is higher, more reciprocity implies lower correlation (Fig. 7f). Networks that solve the s-task but do not minimize the Frobenius norm (*T* networks) showed almost zero reciprocity. This implies that reciprocity is not a property of all networks that solve the s-task. On the contrary, most networks that solve the s-task do not show significant reciprocity, unless other structural constrain, such as Frobenius norm minimization, is imposed. However, Frobenius norm minimization alone only produces negative reciprocity (in random graphs). For positive reciprocity to occur in networks with high number of neurons, both high sequence memory and Frobenius norm minimization is required.

**Figure 7.**
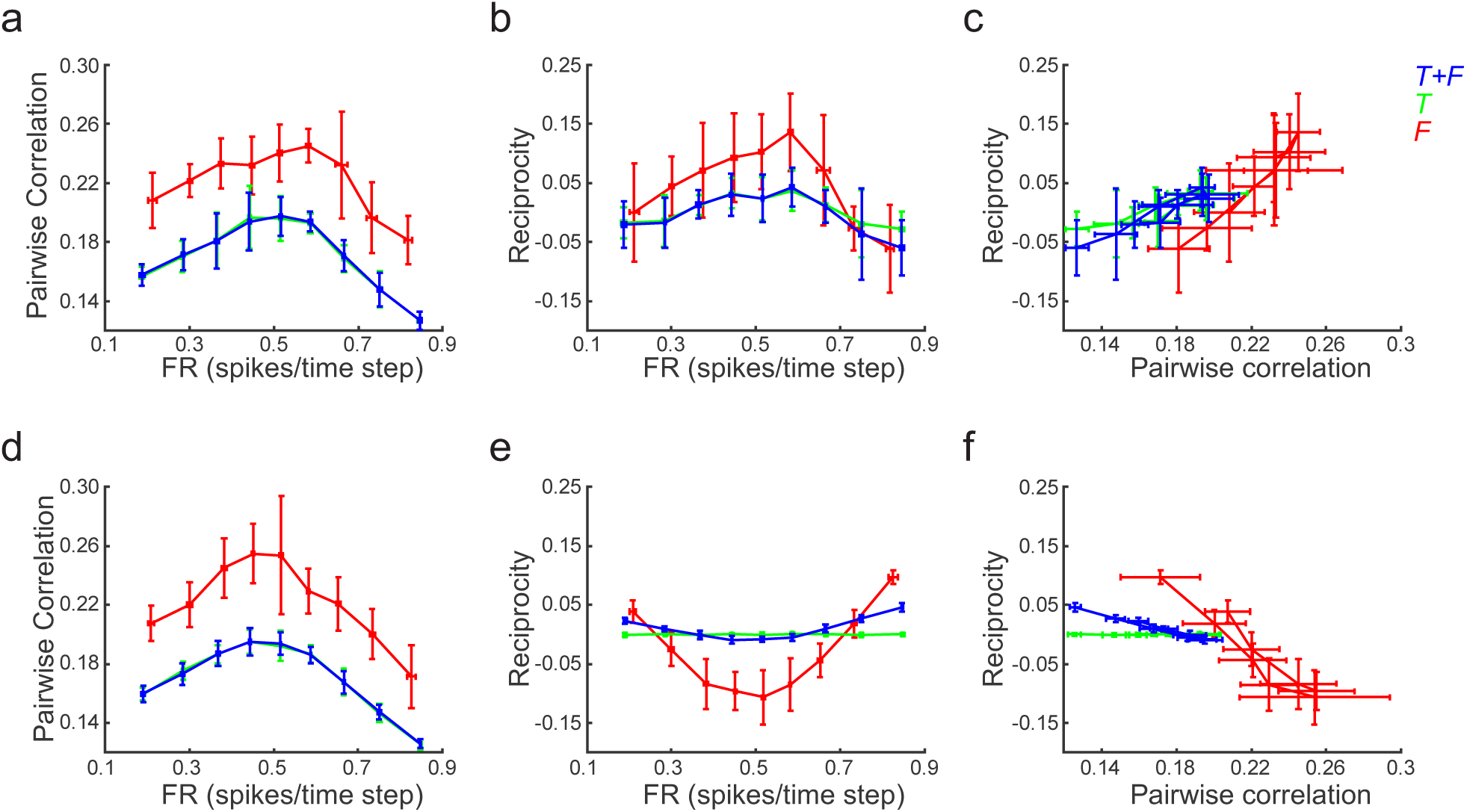
Correlation and reciprocity differentiate networks with sequence memory from random transition networks. **a-c**, networks constructed to solve the s-task with *τ* = 7 and *f*_*r*_ = 1 (*T+F* networks), their isofunction network (*T* network), and networks with the same number of neurons and network states that follow a random transition graph (*F* networks). (a), correlation increases as FR approaches 0.5 spikes/time step. *F* networks show a positive offset with respect to *T+F* and *T* networks. (b), the dependency between reciprocity and FR is similar to the dependency between CC and FR. Higher reciprocity values are found in *F* networks. (c) reciprocity grows linearly with correlation, as expected from panels (a) and (b). **d-f**, idem a-c, but with *f*_*r*_ = 4. (d), the CC/FR relationship is similar to the one observed with lower neuron number (panel (a)). **e**, the reciprocity/FR relationship inverted as the neuron number was increased. Reciprocity is minimized as FR approaches 0.5 spikes/time steps, and increases towards lower or higher FR values. *F* networks show pronounced negative reciprocity. (f) reciprocity decreases linearly with correlation, as expected from panels (d) and (e). Mean ± SD are shown; n = 10 networks were constructed for each target FR and network type. All random graphs were constructed with *f*_*bc*_ = 0.25.

We asked whether the results depicted in Fig. 7 also occur in networks which lack self-connections and comply with Dale’s principle. To that end we imposed these structural constrains to networks constructed with *τ* = 7 and *f*_*r*_ = 4, and found a reciprocity/FR relationship that resembles the one observed in unconstrained networks, with *F* networks showing prominent negative reciprocity and *T+F* networks showing increasing reciprocity as FR departs from 0.5 spikes/time step (Fig. 8a). Correlations increased as FR approached 0.5 spikes/time step, with *F* networks showing more correlation than *T+F* networks (Fig. 8b,c). Correlation in *T+F* and *F* networks were higher for pairs of inhibitory neurons than for pairs of excitatory neurons, as has been observed experimentally^25^, while correlations between excitatory and inhibitory neurons laid in the middle. It is interesting to note that the classification of neurons as excitatory or inhibitory was not defined by design, but emerged during the enforcement of the structural constrains, when the network firing states were already chosen. This suggests that it was the (predefined) firing statistic of the neurons, specially the correlation among them, which ultimately defined which neuron could become excitatory and which inhibitory.

**Figure 8.**
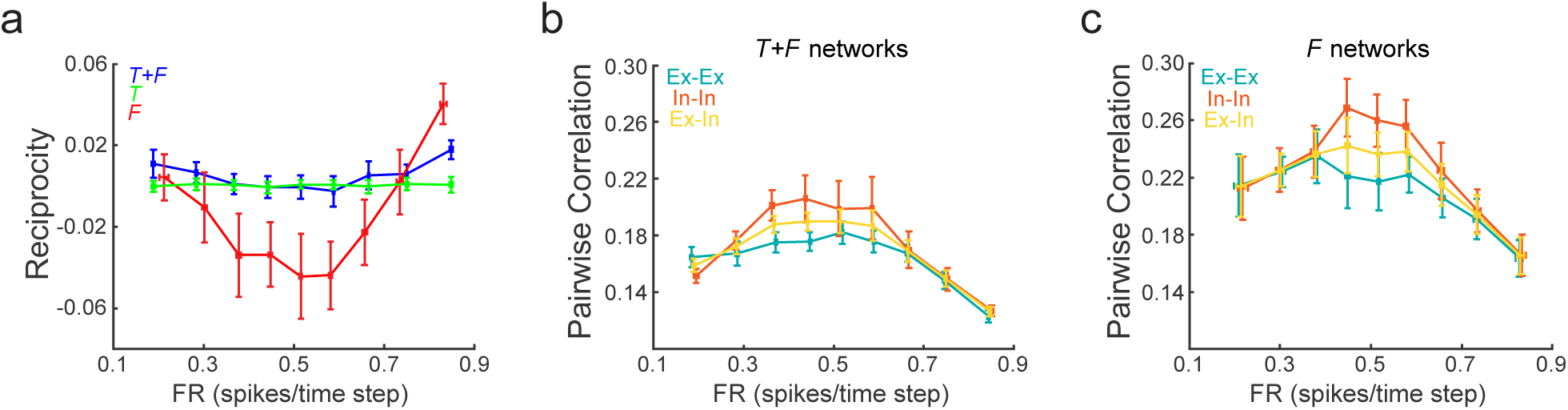
Reciprocity and correlation of structurally constrained networks. **a**, reciprocity as a function of FR for networks without self-connections and Dale’s principle with 1:1 Ex:In ratio. Reciprocity shows a parabolic relationship with FR, decreasing as FR approaches 0.5 spikes/time step. *F* networks show strong negative reciprocity, while *T* networks reciprocity is close to zero. **b**, pairwise correlation for *T+F* networks as a function of firing rate. Correlation was computed over pairs of excitatory neurons (Ex-Ex), pairs of inhibitory neurons (In-In), and pairs of one excitatory and one inhibitory neuron (Ex-In). Correlation has a maximum close to 0.5 spikes/time step. The In-In pairs show the highest correlations, followed by the Ex-In pairs. The Ex-Ex pairs show the lowest correlation. **c**, pairwise correlation for *F* networks as a function of firing rate. The CC/FR relationship is similar to the one observed for *T+F* networks, although *F* networks correlation is displaced towards higher values. Mean ± SD are shown; n = 20 networks were constructed for each target FR and network type. Firing rates of excitatory or inhibitory neurons are displayed for Ex-Ex and In-In curves, respectively. For Ex-In curves the average FR over all neurons is shown.

In the *F* networks studied so far, each network state can be reached from either one of the two stimuli, or from both stimuli. This is the case because the random transition graphs allow nodes with incoming edges from both stimuli. Network states which can be reached from both stimuli (bicoloured nodes, see Methods) codify stimuli in a relative manner, meaning that the identity of the stimulus presented at time step *t* can be decoded if the network state at time step *t* and at time step *t* +1 is known. On the other hand, network states which can be reached from exclusively one of the two stimuli (monocoloured nodes), codify stimuli in an absolute manner, since it is possible to know the identity of the presented stimuli at time step *t* by knowing the network state at time step *t* +1 alone. We asked whether the proportion of relative coding states and absolute coding states could explain the strong differences in correlation and reciprocity found between *T+F* and *F* networks. To that end, we changed the fraction of nominal bicolored nodes *f*_*bc*_ and computed reciprocity for *F* networks of fixed *τ*, FR and *f*_*r*_ (Fig. 9). We found that negative reciprocity values are caused by relative coding network states, since reciprocity is reduced as the fraction of these states is increased. When all network states are absolute coding states (*f*_*bc*_ = 0), reciprocity is the lowest, as observed in *T+F* networks with the same FR and *f*_*r*_. This suggests that reciprocity differentiates networks by how their network states codify stimuli, regardless of the capacity of the network for sequence coding.

**Figure 9.**
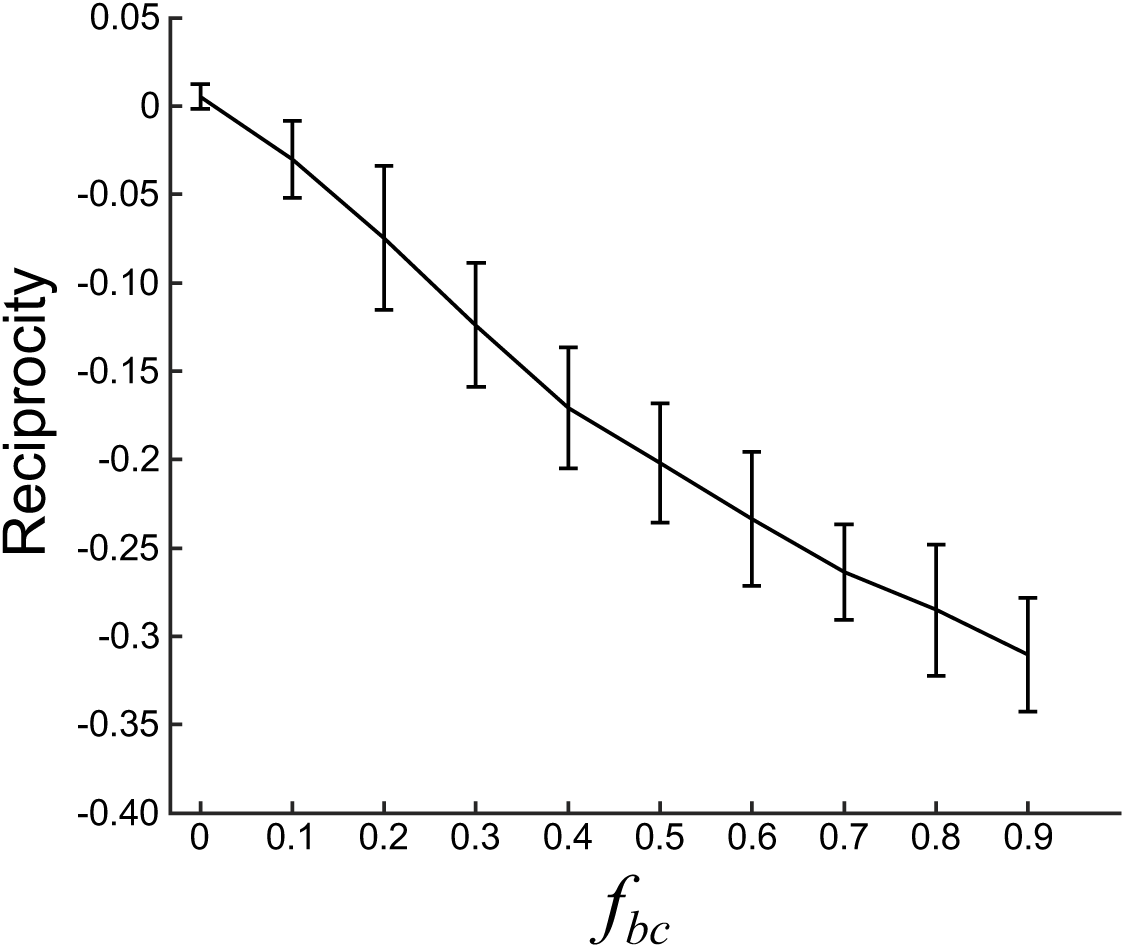
Relative coding network states cause negative reciprocity. Reciprocity as a function of *f*_*bc*_, the fraction of nominal bicolored nodes, in networks that follow random transition graphs. Reciprocity decreases linearly with *f*_*bc*_, approaching zero as *f*_*bc*_ approaches zero. Mean ± SD are shown, n = 30 networks for each *f*_*bc*_. Networks were constructed with target FR = 0.5 spikes/ time step, and with the same number of neurons and network firing states as *T+F* networks constructed with *τ* = 7 and *f*_*r*_ = 4.

## Discussion

We have presented a simple method to generate binary neural network models that accomplish a desire task. Binary networks are computational inexpensive, and despite their simplicity many neurophysiological and neuroanatomical observations have been recapitulated by means of these networks^21,26^. Our key contribution is to note that, for networks in which neurons inputs are linearly added, their synaptic weights can be found by solving a system of linear equations. In turn, this system can be constructed from the transition graph associated with the solution of the target task. System consistency is guarantee if the dependent variables of the system (the neurons activations) are linearly combined following the linear dependences among the independent variables (the firing states). We have shown how the FTP method works with the simplest of networks. Yet, we think the same procedure can be implemented in networks built from more complex neuron models, like the firing rate model or the leaky integrate-and-fire model, provided that a system of linear equations can be constructed.

Current automated methods for constructing network models relay on off-the-shelf optimization algorithms typically employed in the artificial intelligence field, like stochastic gradient descent^3^, genetic algorithms^27^ or evolutionary strategies^28^. These optimization algorithms iteratively change network parameters in a direction that minimizes a loss function, and have proved to be very effective in finding networks that solve very complex tasks^29,30^. However, they require a considerable amount of human design, and there are no guarantees that they can reach a solution. Moreover, each optimization iteration requires the evaluation of the network, which is time consuming, especially for a recurrent network performing in a multi-trial task. In contrast, the FTP algorithm reduces the problem of finding a suitable network to a series of linear combinations and the solution of a linear system, which can be solved in polynomial time. Most importantly, it is guaranteed that the resulting network will solve the task perfectly.

When employing traditional optimization algorithms to fit neural networks, a loss function is defined, taking into account all the required constrains, whether these are task related, activity related, or structural. Then, the loss function is minimized and hence all constrains are enforced at once. In this scenario the relationship between parameters and the loss function can be quite complex, and some conflict between constrains may emerge. Conversely, one key advantage of our method is that it allows to uncouple the dynamic and coding aspects of the network from the structural aspects, giving the opportunity of sampling them independently. Since the method proceeds from the firing states to the parameters, it allows to find networks with desired activity profiles, and to study the resulting connectivity. Further structural constrains can be enforced in a second stage, by projected gradient descent, or any other optimization algorithm. The fact that projected gradient descent worked so well suggests that structural constrains are easy to implement once the connectivity required to solve the task is in place. This is probably because structural measures such as sparsity, clustering, etc are a simple function of the synaptic weights. On the contrary, the relationship between weights and network function or coding capabilities are more complex. In this way, our method gives more control over each constrain type, making the whole process simpler at the same time.

To show the applicability of the method we employed it to construct networks that solve a stimuli sequence memory task (s-task) in which the network has to codify in its network firing states the sequence of the last *τ* stimuli that were presented. This task is relevant in the broad sense of working memory function. Although working memory is traditionally associated with maintaining information about a single stimulus in the persistent activity of recurrently connected neurons^31,32^, mounting evidence suggests that neuron populations code information in the form of highly heterogenous firing sequences^33,34^. Sustained activity can be a suitable strategy when there is one specific relevant stimulus to attend, whose identity has been already elucidated. However, more complex scenarios require keeping track of sequences of stimuli. An example of this case is the processing of language, in which the succession of utterances must be integrated over time, from phonemes, to words, to phrases, so that the meaning of speech depends on the whole sequence^35^. We explored the case of two stimuli presented with equal probability, but the analysis could be extended to more realistic cases in which stimuli presentation probability is not uniform. It is expected that statistical regularities in the input sequences are going to be exploited by the network, resulting in more specialized connectivities. The relationship between sequence statistics and network structure should be further studied. For example, it would be of interest the case in which each stimulus lasts more than one time-step, and they are interleaved by neutral stimuli, which could act as distractors. Then, the relationship between feedback and feedforward connections could be studied, in relation with the duration of each stimulus presentation, and with that of the distractors.

The structure-function relationship is central to neuroscience^36–38^. Connectivity at the macro meso and micro scale, neurons biophysics, plasticity mechanisms, among other structural traits, all act co-ordinately to give sophisticated adaptive behaviour. It is widely believed that structural properties of networks have evolved over time to proficiently perform function, many times in an optimal way^39,40^. However, brain structure could also be the result of other constrains, different from those imposed by adaptive behaviour. For example, neural network modularity might have emerged as a good structural trait for solving tasks which have a modular or hierarchical aspect^41^. But it could also have emerged as a result of previously acquired structural traits such as constrains in the length of dendrites and axons, which precludes the possibility of a much wider connectivity. Thus, determining how much of the structure observed in the brain comes from task-related constrains and how much comes from other structural traits is central to understanding the structure/function relationship. A theoretical approximation to this issue consists in constructing neural network models that solve different kind of tasks under a variety of structural constrains, and then study the pattern of connectivity that emerges and relate it to the experimentally observed connectivity in real brains. This approximation requires to sample as uniformly as possible from the set of networks that fit the task and structural constrains. However, optimization methods commonly used in network parameter fitting may give a restricted set of solutions, thus biasing any conclusion about the structure/function relationship. Another issue is that some connectivity traits could emerge only in networks of certain size, and fitted to several tasks. In this case, fitting large networks with complex cost functions could have a high computational cost. Consequently, generating a relatively large sample of networks suitable for statistical treatment of their connectivity would result unfeasible. In this aspect, the FTP algorithm is very well suited for answering structure/function questions, since exact solutions can be computed starting from an arbitrary set of population firing codes, as long as it defines a system of equations that have a solution.

The FTP approach allows to test hypothesis linking structure and function by constructing networks which follow transition graphs that instantiate some null hypothesis. Following this approach, we constructed networks which had the same number of network states and neurons required to solve the s-task, but whose state transitions were chosen at random. With this tool at hand we were able to show that a structural feature emerges as *τ* and redundancy increase, evidenced in the reciprocity of the network. The same procedure can be followed to build any other set of networks in accordance with some relevant null hypothesis. Such networks can be easily constructed with the FTP method, while they would be hard to construct with regular optimization algorithms.

Evidence for high reciprocity has been found experimentally, by measuring excitatory postsynaptic potentials of reciprocally connected neurons in vitro^20^. It has also been the centre of theoretical analysis. For example, it has been shown that high reciprocity is recapitulated in networks of binary neurons that have maximum number of attractors^21^. Interestingly, the same work shows that reciprocity is lost when networks are optimized to remember sequences of uncorrelated network states. However, we showed that, when networks are built to codify sequences of stimuli, the network itself shows sequences of states that follow the sequences of stimuli up to an arbitrary *τ*.

Reciprocity was absent in networks taken at random from the set of all connectivities that give the same dynamics. This implies that the observed reciprocity is the result of following a particular transition graph with the additional constrain of weights minimizing the Frobenius norm, the latter being explained biologically as an upper bound on the size of the synapses. Thus, to explain one structural feature (reciprocity), a functional feature (solving the s-task) and another structural feature (Frobenius norm minimization) were required. It would be interesting to study to what extent other structural features encountered in biological neural networks, like modularity or sparsity of connections, can be explained as the answer to some computational demand of adaptive behaviour, or they are the result of another structural feature, or both factors interact, as is the case of the s-task.

In conclusion, we have provided a method that inverts the usual process of constructing neural network models. It allows to probe the dependency between the firing statistics, connectivity and function of a network in a way that is not matched by current optimization algorithms. Moreover, it is computationally inexpensive. Therefore, we consider the method to be a powerful alternative to the way neural networks are constructed to model brain function.

## Methods

### Network simulation and synaptic weights statistics

Networks were evaluated in the s-task during at least *N* _*iter*_ = 10.2^*τ*^ time steps, to gather enough samples of each network state. To assess the similarity between the synaptic weight distribution and a normal distribution we computed the Kolmogorov-Smirnov two samples statistic, between the set of synaptic weights and a set of normally distributed values of the same mean, variance and sample size than that of the synaptic weights.

Equation (4) gives the matrix *w* with lowest Frobenius norm^10^. Since we are considering networks with *N* = 2^*τ*^ + 2 total neurons (including sensory and integration neurons), there are infinite solution matrices *w* for the same system of equations defined by *C* and *U*. These solutions lay in a subspace of ℝ^*N*^ of dimension *N* − rank(*C*). The set of all solutions can be, obtained by finding a matrix Δ*w* such that:

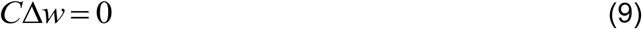

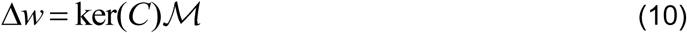

where ker(*C*) is an orthonormal basis of the null space of *C* of dimensions *N* x(*N*_*z*_ − rank(*C*)), 0 is a matrix of zeros, and *M* is a linear mapping of dimensions (*N*_*z*_ − rank(*C*))x *N*_*z*_. These networks share the same stimulus-response mapping. We say they conform an isofunction space.

For several applications, networks with *N* _*z*_ = 2^*τ*^ were desired. Hence, we defined *N* _*z*_ = 2^*τ*^ *f*_*r*_, where *f*_*r*_ stands for ‘redundancy factor’, as the network has *f*_*r*_ - times more neurons than required to solve the s-task with that specific *τ*.

### Computation of fitting times for FTP and a genetic algorithm

We assessed the efficiency of the FTP algorithm by measuring the time expended in finding networks that solve an s-task with *τ* = 1 to *τ* = 12, and *f*_*r*_ = 1.

We also computed the time expended by a genetic algorithm (GA) to obtain networks that solve the s-task of *τ* = 1 to *τ* = 10. We employed a population of *N*_*pob*_ = 200 individuals, each one composed by one matrix *w*_*y*_, one matrix *w*_*r*_ and one vector *θ*. Networks were evaluated in the s-task for *N* _*iter*_ = 10.2^*τ*^ time steps, and its fitness *F* was defined as 1 minus the classification loss of a support vector machine, trained to classify the stimulus presented at time step *t* −*τ* +1 based on the population firing state at time step *t*. The classification model was cross-validated with the holdout method, trained on 50% of the data and tested on the remaining 50%. We picked *T* = 0.1*N*_*pob*_ individuals with the highest fitness as parents. Then, we picked parents at random and built the next generation by mutating each synaptic weight with gaussian noise of zero mean and standard deviation *f*_*mut*_ = 0.1*σ*(*w*). The factor *σ*(*w*) is the standard deviation of the synaptic weights in *w*_*y*_ of networks generated with FTP if we are mutating *w*_*y*_, and the standard deviation of synaptic weights in *w*_*r*_ if we are mutating *w*_*r*_. We defined *f*_*mut*_ in this way to avoid *f*_*mut*_ values that are so big that a solution can not be reached, or so small that the solution will not be reached in a reasonable amount of time. One of the individuals of each generation was an unmutated copy of the best individual of the previous generation (elitism). Threshold vectors *θ* were not mutated. The GA was run until the average fitness surpassed *F*_*target*_ = 0.9, or after 60 minutes of search. Once the stopping criterion was met, the elite individual was evaluated during 100.2^*τ*^ time steps to obtain the final fitness.

We also tested how stable was a solution obtained with FTP under evolution with a GA. We employed FTP to construct networks of given *τ* and *f*_*r*_ = 1. From this network a population of *N*_*pob*_ = 100 individuals was constructed by mutating the network with a mutation rate 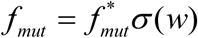, where *σ*(*w*) is the standard deviation of the synaptic weights obtained with FTP. As before, *w*_*y*_ and *w*_*r*_ has their own *f*_*mut*_, according to their corresponding standard deviation. Then, a GA was employed to reduce fitness during 20 generations, and then to increment it for another 20 generations, employing the same 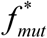. We followed the procedure for *τ* ranging from *τ* = 1 to *τ* = 6, and 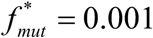 to 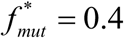 in steps of 0.1. We performed 10 repetitions for each *τ* and 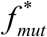. combination.

### Imposing activity constrains

To construct networks with desired FR we generated *U*_*base*_ as described in the Results Section, but adjusted the sign of *r*(*m,i*) such that, after thresholding, matrix *C* had a fraction of ones equals the target FR. To induce signal correlation, we scaled vector Δ by a factor *f*_*cc*_. This manipulation makes each neuron to have very different firing rates for *s*_1_ and *s*_2_, which increases the signal correlation. By following this procedure, we constructed networks in Fig. 4. The target FR values were taken from the range between 0.1 to 0.9 spikes/time step, in steps of 0.1 spikes/time step. The *f*_*cc*_ values were taken from the range between 1 and 10 in unitary steps. A total of 30 networks were constructed for each combination of FR and *f*_*cc*_ values within those ranges. For each network the average FR was computed over the FR of all neurons in the network. Similarly, the average correlation coefficient (CC) was computed from the Spearman correlation coefficient computed for all neuron pairs.

We also employed a GA to evolve a population of *U*_*base*_ matrices to fit their mean FR and correlation. We employed a population of *N*_*pob*_ = 200 individuals, each one composed of one matrix *U*_*base*_ and one vector *θ*. For each individual we constructed *U* and *C* matrices, and computed an approximate value of FR and correlation, under the assumption that each network firing state occurs with equal probability. The fitness *F* of an individual was computed as

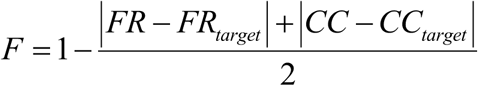

where *FR* and *CC* are the firing rate and correlation values of the networks output, and *FR*_*target*_ and *Cc*_*target*_ are the firing rate and correlation values we want the networks to have.

If an individual produced an inconsistent system, or a system with not enough network states, its fitness was set to zero. We chose *T* = 0.1*N*_*pob*_ and mutated *U*_*base*_ by adding gaussian noise to each matrix element, of zero mean and standard deviation *σ* = *f*_*mut*_ = 0.1. Threshold vectors *θ* were not mutated. Elitism was employed. The GA was run until the average fitness surpassed *F*_*target*_ = 0.95. Firing rates and correlations shown in Fig. 4d were computed by running the network constructed from the elite *U*_*base*_ during 30.2^*τ*^ time steps.

### Imposing structural constrains

Solving equation (4) gives networks with minimum Frobenius norm. These networks do not suffice basic structural features observed experimentally, such as the lack of self-connections or Dale’s principle. To impose such structural constrains we constructed a matrix Δ*w*_*d*_ such that *w*_*d*_ = *w* + Δ*w*_*d*_. Matrix *w*_*d*_ is a matrix which fulfils the desired structural constrains. Most probably Δ*w*_*d*_ will not be within the null space of *C*, and thus *w*_*d*_ will not be a solution to the system defined by *C* and *U*. Hence, we defined a matrix:

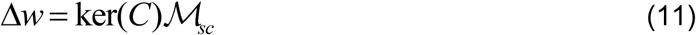

Where *ℳ*_*sc*_ = ker(*C*)^+^ Δ*w*_*d*_, and ker(*C*)^+^ is the Moore-Penrose pseudoinverse of ker(*C*). Matrix *ℳ*_*sc*_ is a linear mapping that incorporates the desired structural constrains, making Δ*w* the change in matrix *w* within the null space of *C* that is closest to Δ*w*_*d*_, in the least squares sense.

We imposed three structural constrains: no self-connections, Dale’s principle, and a certain degree of sparsity. Thus:

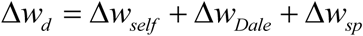

The matrix Δ*w*_*self*_ (which deletes self-connections in the integration neurons) has values:

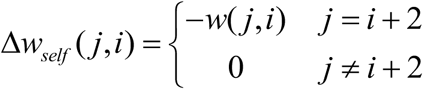

Matrix Δ*w*_*dale*_ was defined as:

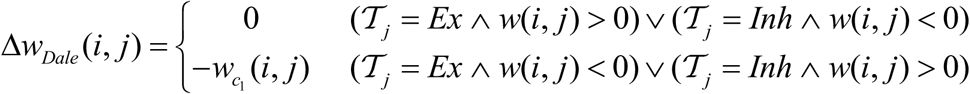

Where 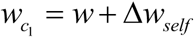 and *T*_*j*_ ∈{*Ex, In*} indicates if neuron *j* was chosen to be excitatory (*Ex*) or inhibitory (*Inh*). Matrix Δ*w*_*Dale*_ sets to zero the synaptic weights that violate Dale’s principle. Neuron *j* was chosen to be excitatory if 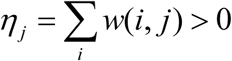. Otherwise, it was chosen to be inhibitory. If more excitatory/inhibitory neurons were required, neurons with negative/positive *η* closest to 0 were set as excitatory/inhibitory as needed.

Matrix Δ*w*_*sp*_ to enforce sparsity was defined as:

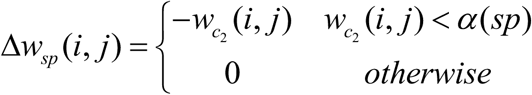

Where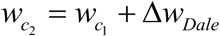. The value *α* (*sp*) is the *sp* percentile of the absolute values in 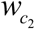. In this manner Δ*w*_*sp*_ will set to zero the lowest weights such that a sparsity *sp* is enforced.

The loss function ℒ(*k*) at iteration *k* was defined as the average of the absolute Δ*w*_*d*_ (*i, j*) values:

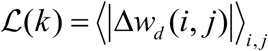

Where ⟨ ⟩_*i, j*_ stands for the average across indexes (*i, j*). The structural constrains were imposed through an iterative process, in which at each iteration the neurons were classified as excitatory or inhibitory according to their *η*, a matrix Δ*w* was computed using eq. (11), and a new *w* was obtained. The process was stopped when the loss fell below a desired value *e*_1_, in which case the fitting process was considered successful. The process was also stopped if (*L*(*k*) − *L*(*k* −1)) / *L*(*k*) < *e*_2_. When this latter condition was met, the fitting process was considered unsuccessful, since the error was not decreasing fast enough and would probably converge to an unacceptable value above zero. We used *e*_1_ = 10^−3^ and *e*_2_ = 10^−4^. If the process was successful, values that violated any of the constrains were clipped to zero. These values are expected to be small enough since the error was small. We computed:

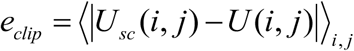

Where *U*_*sc*_ = *Cw*_*sc*_ with *w*_*sc*_ being the resulting synaptic weight matrix after the constraining process, to verify that the deviation from the original *U* was negligible. If the process was unsuccessful, or the clipping error *e*_*clip*_ > 10^−3^, then the original *w* was considered not to be suitable for the structural fitting.

We measured the efficiency of the process by computing the number of networks generated (*# attempts*) and the running time *t* expended until reaching the first successfully constrained network. We varied *τ* from *τ* = 3 to *τ* = 6. For each *τ* we varied the number of integration neurons in steps of 16 neurons, from a minimum number of 4.2^*τ*^ to the maximum value 2^6^. For each combination of *τ* and neuron number we generated networks with the FTP algorithm, and subjected them to structural constrain (no self-connections, 4:1 Ex:In ratio, and a minimum sparsity *sp* = 40%). We obtained 10 measurements of *# attempts* and *t*, from which we computed the mean and SD depicted in Fig. 5e,f.

### Network construction from random transition graphs

To construct random transition graphs that have an associated consistent system, we first constructed a matrix *U* _*base*_, a vector Δ, and a matrix *U*^*^ of *M* = 2^*τ*^ rows such that the row vectors 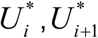 and Δ were linear combinations, with indexes *i* being odd numbers between 1 and *M*. Next, we constructed matrix *U* in a way that ensures that each node in the graph was reachable, meaning that every node had to receive at least one edge. This is equivalent to say that every row in *U*^*^ is found at least once in *U*. Therefore, we set the first *M* rows in *U* equal to the *U*^*^ matrix. Then, to define the remaining rows in *U*, we chose *M* /2 pairs of indexes *i,i* +1, picking *i* values at random from the set of odd numbers between 1 and *M*. In this way, and unlike the transition graphs that solve an s-task, nodes could receive just one edge, or more than 2.

So far, if a row vector 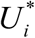 appeared in matrix *U* more than once, then it appeared only in odd rows, or only in even rows, but not in both. This is because, indexes were ordered from 1 to *M* in the first half of *U*, and ordered in pairs of *i,i* +1 indexes in the second half. The resulting graph would be one in which any given node is reachable as the result of the presentation of either *s*_1_ or *s*_2_, but not from both. In other words, if node *b* is reachable from node *a* after *s*_*i*_ presentation, then node *b* is reachable from node *c* only after *s*_*i*_ presentation, where *c* is any other node from which *b* is reachable. Following the colour code of the graph in Fig. 1b, any node receives arrows of the same colour. We wanted graphs as random as possible, so nodes reachable thought different stimuli were desired. We define these nodes as *bicoloured* nodes. In terms of indexes in matrix *U*, a bicolored node translates into a row vector 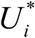 that appeared in matrix *U* in both odd and even row. For example, if we had indexes (1, 2,3, 4) for the first 4 rows of *U*, with (*U*_1_,*U*_2_, Δ) and (*U*_3_, *U*_4_, Δ) each being linearly combined, then we wanted to change this series to (1, 2, 2, 4), or (1, 2,3,1). This requires to generate new linear combinations, in particular, (*U*_1_,*U*_2_, *U*_4_, Δ) will be linearly combined, for the first example, and (*U*_1_,*U*_2_, *U*_3_, Δ) in the second example. Thus, we modified matrix *U* to generate *f*_*bc*_*M* /4 bicolored nodes, where *f*^*bc*^ stands for ‘bicolored fraction’ and is a number between 0 and 1. The (nominal) maximum number of bicolored nodes is *M* / 4, since we generated one node for each series of indexes *i* to *i* + 3. Finally, we constructed matrix *Z* by thresholding matrix *U*^*^, and then matrix *C*, which rows were in the form:

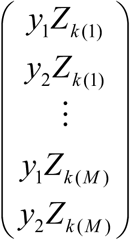

where *Z*_*i*_ is the ith row vector in matrix *Z* and vector *k* is a permutation of the list of integers from 1 to *M*.

Following the above procedure, we constructed random transition graphs that respected the linear combinations needed so that a consistent system of equations could be constructed. Given that these networks do not solve the s-task but are the minimum Frobenius norm solution to a random transition graph, we call them *F* networks. The procedure avoids index sequences like (1221), since this ordering gives a consistent system only if Δ= 0, in which case stimuli cannot be distinguished by the network. The procedure also avoids index sequences of the type (11) (one node leads to another single node through both stimuli, *s*_1_ and *s*_2_). If this were the case, one possibility is that *s*_1_ and *s*_2_ produce the same *u* values. Therefore, Δ is a vector of zeros and stimuli cannot be discriminated. Another possibility is that stimuli lead to different vectors *u*, but these vectors in turn lead to the same vector *c* after thresholding. This situation is possible, but would require careful selection of *u* values in relation to *θ*, and for this it was avoided.

We constructed networks that follow random transition graphs and compared their properties with the properties of networks that solve the s-task. In particular, we measured the reciprocity of the network, defined as the Spearman correlation between weights of incoming and outgoing synapses. Reciprocity was computed over matrix *ω*, a normalized version of the synaptic weights constructed by taking the absolute values of *w* and scaling them between 0 and 1. Since imposing structural constrains like Dale’s principle, or sparsity, may generate many zero-valued weights, reciprocity was computed over *ω* values such that *ω* (*i, j*) ≠ 0 and *ω* (*i, j*) ≠ 0 for each possible pair (*i, j*).

For each network generated to solve the s-task we also picked a network from its isofunction space, that is, from the set of all networks that has the same stimulus-response mapping (as described above). We refer to these networks as *T* networks, since they solve the s-task but they are not the minimum Frobenius norm solution. The linear mapping ℳ in eq. (10) has entries 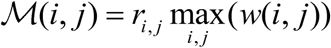, where *r*_*i, j*_ is a random number, different for each entry, sampled uniformly from the [−1,1] interval, and *w* is the synaptic weight matrix from which an isofuntion network is desired. We defined mapping ℳ in this way to obtain isofunction networks with synaptic weight values within the range of the weights in the original network.

In addition, for each network that solves the s-task we constructed an isofunction network with structural constrains: no self-connections and Dale’s principle with excitatory and inhibitory neurons in equal numbers.

All the algorithms were implemented in Matlab.

## Acknowledgements

This research was supported by grants from Agencia Nacional de Promoción Científica y Tecnológica (ANPCYP) PICT 2016-2145 and PICT 2017-3208) and from Universidad de Buenos Aires (UBACYT 20020170100568BA). The work of Camilo J. Mininni was supported by a postdoctoral fellowship from ANPCYT.

## Author contributions

CJM conceived the project, implemented the algorithms and wrote the initial draft. CJM and BSZ reviewed and edited the final manuscript.

## Materials and correspondence

Correspondence to Camilo J. Mininni.

Matlab codes are available upon request.

## References

1. Kriegeskorte, N. & Douglas, P. K. Cognitive computational neuroscience. Nat. Neurosci. 21, 1148–1160 (2018).

2. Bassett, D. S., Zurn, P. & Gold, J. I. On the nature and use of models in network neuroscience. Nat. Rev. Neurosci. 19, 566–578 (2018).

3. Song, H. F., Yang, G. R. & Wang, X. J. Training Excitatory-Inhibitory Recurrent Neural Networks for Cognitive Tasks: A Simple and Flexible Framework. PLoS Comput. Biol. 12, 1–30 (2016).

4. Richards, B. A. et al. A deep learning framework for neuroscience. Nat. Neurosci. 22, 1761–1770 (2019).

5. Barrett, D. G., Morcos, A. S. & Macke, J. H. Analyzing biological and artificial neural networks: challenges with opportunities for synergy? Curr. Opin. Neurobiol. 55, 55–64 (2019).

6. Bengio, Y. & LeCun, Y. Scaling Learning Algorithms Towards AI. in Large-Scale Kernel Machines (eds. Bottou, L., Chapelle, O., DeCoste, D. & Weston, J.) 321–358 (The MIT Press, 2007).

7. Sussillo, D., Churchland, M. M., Kaufman, M. T. & Shenoy, K. V. A neural network that finds a naturalistic solution for the production of muscle activity. Nat. Neurosci. 18, 1025–1033 (2015).

8. Maheswaranathan, N., Williams, A. H., Golub, M. D., Ganguli, S. & Sussillo, D. Universality and individuality in neural dynamics across large populations of recurrent networks. (2019).

9. Penrose, R. & Todd, J. A. On best approximate solutions of linear matrix equations. Math. Proc. Cambridge Philos. Soc. 52, 17–19 (1956).

10. Neri, F. *Linear algebra for computational sciences and engineering*. Linear Algebra for Computational Sciences and Engineering (Springer, 2016). doi:10.1007/978-3-319-40341-0

11. Lennie, P. & Place, W. The Cost of Cortical Computation. Curr. Biol. 13, 493–497 (2003).

12. Mizuseki, K. & Buzsáki, G. Preconfigured, Skewed Distribution of Firing Rates in the Hippocampus and Entorhinal Cortex. Cell Rep. 4, 1010–1021 (2013).

13. Schneidman, E., Berry, M. J., Segev, R. & Bialek, W. Weak pairwise correlations imply strongly correlated network states in a neural population. Nature 440, 1007–1012 (2006).

14. Strata, P. & Harvey, R. Dale’s principle. Brain Res. Bull. 50, 349–350 (1999).

15. Lefort, S., Tomm, C., Floyd Sarria, J. C. & Petersen, C. C. H. The Excitatory Neuronal Network of the C2 Barrel Column in Mouse Primary Somatosensory Cortex. Neuron 61, 301–316 (2009).

16. Seeman, S. C. et al. Sparse recurrent excitatory connectivity in the microcircuit of the adult mouse and human cortex. Elife 7, 1–27 (2018).

17. Szegedi, V. et al. Robust perisomatic GABAergic selfinnervation inhibits basket cells in the human and mouse supragranular neocortex. Elife 9, 1–19 (2020).

18. Bekkers, J. M. Neurophysiology: Are autapses prodigal synapses? Curr. Biol. 8, 52–55 (1998).

19. Calamai, P. H. & Moré, J. J. Projected gradient methods for linearly constrained problems. Math. Program. 39, 93–116 (1987).

20. Song, S., Sjöström, P. J., Reigl, M., Nelson, S. & Chklovskii, D. B. Highly Nonrandom Features of Synaptic Connectivity in Local Cortical Circuits. PLoS Biol. 3, e68 (2005).

21. Brunel, N. Is cortical connectivity optimized for storing information? Nat. Neurosci. 19, 749–755 (2016).

22. Garlaschelli, D. & Loffredo, M. I. Patterns of link reciprocity in directed networks. Phys. Rev. Lett. 93, 1–4 (2004).

23. Schulz, D. P. A., Sahani, M. & Carandini, M. Five key factors determining pairwise correlations in visual cortex. J. Neurophysiol. 114, 1022–1033 (2015).

24. de la Rocha, J., Doiron, B., Shea-Brown, E., Josić, K. & Reyes, A. Correlation between neural spike trains increases with firing rate. Nature 448, 802–806 (2007).

25. Constantinidis, C. & Goldman-Rakic, P. S. Correlated discharges among putative pyramidal neurons and interneurons in the primate prefrontal cortex. J. Neurophysiol. 88, 3487–3497 (2002).

26. Alemi, A., Baldassi, C., Brunel, N. & Zecchina, R. A Three-Threshold Learning Rule Approaches the Maximal Capacity of Recurrent Neural Networks. PLoS Comput. Biol. 11, 1–23 (2015).

27. Such, F. P. et al. Deep Neuroevolution: Genetic Algorithms Are a Competitive Alternative for Training Deep Neural Networks for Reinforcement Learning. (2017). doi:1712.06567

28. Salimans. Evolution Strategies as a Scalable Alternative to Reinforcement Learning. Proc. - 4th IEEE Int. Conf. Softw. Testing, Verif. Valid. Work. ICSTW 2011 476–485 (2011). doi:10.1109/ICSTW.2011.58

29. Vinyals, O. et al. Grandmaster level in StarCraft II using multi-agent reinforcement learning. Nature 575, 350–354 (2019).

30. Silver, D. et al. A general reinforcement learning algorithm that masters chess, shogi, and Go through self-play. Science (80-.). 362, 1140–1144 (2018).

31. Kojima, S. & Goldman-Rakic, P. S. Delay-related activity of prefrontal neurons in rhesus monkeys performing delayed response. Brain Res. 248, 43–50 (1982).

32. Guo, Z. V. et al. Maintenance of persistent activity in a frontal thalamocortical loop. Nature 545, 181–186 (2017).

33. Orhan, A. E. & Ma, W. J. A diverse range of factors affect the nature of neural representations underlying short-term memory. Nat. Neurosci. 22, 275–283 (2019).

34. Murray, J. D. et al. Stable population coding for working memory coexists with heterogeneous neural dynamics in prefrontal cortex. Proc. Natl. Acad. Sci. U. S. A. 114, 394–399 (2017).

35. Vries, M. H. De, Christiansen, M. H. & Petersson, K. M. Learning Recursion: Multiple Nested and Crossed Dependencies. Biolinguistics 10–35 (2011).

36. Honey, C. J., Thivierge, J. P. & Sporns, O. Can structure predict function in the human brain? Neuroimage 52, 766–776 (2010).

37. Bullmore, E. & Sporns, O. Complex brain networks: Graph theoretical analysis of structural and functional systems. Nat. Rev. Neurosci. 10, 186–198 (2009).

38. Vázquez-Rodríguez, B. et al. Gradients of structure–function tethering across neocortex. Proc. Natl. Acad. Sci. U. S. A. 116, 21219–21227 (2019).

39. Kording, K. P. Bayesian statistics: Relevant for the brain? Curr. Opin. Neurobiol. 25, 130–133 (2014).

40. Knill, D. C. & Pouget, A. The Bayesian brain: the role of uncertainty in neural coding and computation. Trends Neurosci. 27, (2004).

41. Ellefsen, K. O., Mouret, J. B. & Clune, J. Neural Modularity Helps Organisms Evolve to Learn New Skills without Forgetting Old Skills. PLoS Comput. Biol. 11, 1–24 (2015).

